# An integrative analysis of cell-specific transcriptomics and nuclear proteomics of sleep-deprived mouse cerebral cortex

**DOI:** 10.1101/2024.09.24.611806

**Authors:** Pawan K. Jha, Utham K. Valekunja, Akhilesh B. Reddy

**Affiliations:** Department of Systems Pharmacology & Translational Therapeutics, Perelman School of Medicine, University of Pennsylvania, Philadelphia, PA 19104, USA; Institute for Translational Medicine and Therapeutics, Perelman School of Medicine, University of Pennsylvania, Philadelphia, PA 19104, USA; Chronobiology and Sleep Institute (CSI), Perelman School of Medicine, University of Pennsylvania, Philadelphia, PA 19104, USA

## Abstract

Sleep regulation follows a homeostatic pattern. The mammalian cerebral cortex is the repository of homeostatic sleep drive and neurons and astrocytes of the cortex are principal responders of sleep need. The molecular mechanisms by which these two cell types respond to sleep loss are not yet clearly understood. By combining cell-type specific transcriptomics and nuclear proteomics we investigated how sleep loss affects the cellular composition and molecular profiles of these two cell types in a focused approach. The results indicate that sleep deprivation regulates gene expression and nuclear protein abundance in a cell-type-specific manner. Our integrated multi-omics analysis suggests that this distinction arises because neurons and astrocytes employ different gene regulatory strategies under accumulated sleep pressure. These findings provide a comprehensive view of the effects of sleep deprivation on gene regulation in neurons and astrocytes.

## Introduction

Sleep is ubiquitous in species with well-developed neural networks and is essential for maintaining brain and body physiology ^1–4^. Sleep loss impairs cognition, memory, metabolism, and immune function ^5–8^. However, the molecular basis of sleep and the mechanisms underlying the deleterious effects of sleep deprivation remain largely unknown.

The mammalian cerebral cortex is a key regulator of homeostatic sleep drive ^9,10^. Cortical astrocytes and neurons drive crucial homeostatic brain functions ^11,12^. We previously showed that sleep deprivation modulates astrocyte-neuron signaling and differentially alters transcription, translation, and post-translational profiles in these cell types ^13^. Recent single-cell or nuclear transcriptomics studies of various brain regions under different sleep pressures corroborate our findings that sleep need regulates gene expression in a cell-specific manner ^13–15^. However, transcriptomics data may not accurately reflect functional protein abundance ^16,17^, as some proteins may reside in, or traffic to, the nucleus to orchestrate gene expression programs.

To investigate how sleep need modulates cellular processes and gene regulation in a cell-type-specific manner, we performed simultaneous transcriptional and global nuclear proteome profiling of cortical neurons and astrocytes from sleep-deprived mice. Integrated analysis of all readouts from both cell types revealed distinct patterns of gene expression, nuclear protein abundance, and putative regulatory factor networks driving the transcriptional response to sleep deprivation. This study thus provides insights into the gene regulation strategies employed by neurons and astrocytes in response to sleep loss and serves as a resource for future investigations of cell-type-specific mechanisms of sleep regulation.

## Results

### Sleep deprivation alters transcriptional profiles of neurons and astrocytes in the cortex

To study how sleep loss impacts the transcriptional profiles of neurons and astrocytes, we sleep-deprived (SD) a group of mice in their usual resting period, from ZT0 to ZT12 (ZT; *zeitgeber time*). A control group was allowed to sleep *ad libitum* (normal sleep, NS). At ZT12, we collected the brain and dissected the cortex for cell suspension preparation, then separated neurons and astrocytes for transcriptomics and nuclear proteomics ^13^ (Fig. 1 A-B).

**FIGURE 1.**
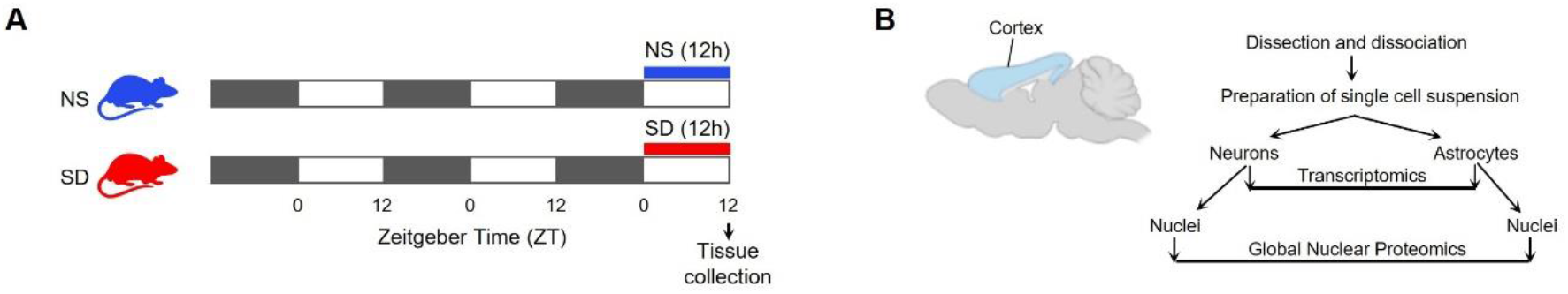
Schematic representation of experimental strategy. (**A**) sleep experiment and tissue collection timeline. Black bars represent darkness, white bars represent periods of light, and top blue and red bars represent normal sleep (NS), and sleep deprivation (SD). (**B**) Workflow for transcriptomics and nuclear proteomics. Cortex of the brain, illustrated by a light blue shade, was dissected for astrocyte and neuron separations. Cells were used for transcriptomics and global nuclear proteomics.

Sleep deprivation significantly altered the expression of 6.18% (1804/29170) and 1.32% (357/27046) of genes in neurons and astrocytes, respectively (FDR < 0.05). Applying a 30% fold-change threshold (1.3-fold cutoff, FDR < 0.05), 547 genes in neurons and 231 in astrocytes were differentially expressed, with the majority downregulated in neurons and upregulated in astrocytes (Fig. 2A-B, Fig. 3A-B). Of these, 18 genes exhibited the same directionality in both cell types, while 5 showed opposite trends (Table 1). Of note, several alterations such as in *Cdkn1a* (Cyclin-dependent kinase inhibitor 1), *Il1r1* (Interleukin-1 receptor type 1), *Map3k6* (Mitogen-activated protein kinase kinase kinase 6), *Syt12* (Synaptotagmin-12), *Npas4* (Neuronal PAS domain-containing protein 4), *Ptgs1* (Prostaglandin G/H synthase 1), *Hspa1a* (Heat shock 70 kDa protein 1A), *Hspa1b* (Heat shock 70 kDa protein 1B), *Dbp* (D site-binding protein) and *Phyhd1* (Phytanoyl-CoA dioxygenase domain-containing protein 1) were previously reported in sleep studies ^13,15,18–21^.

**FIGURE 2.**
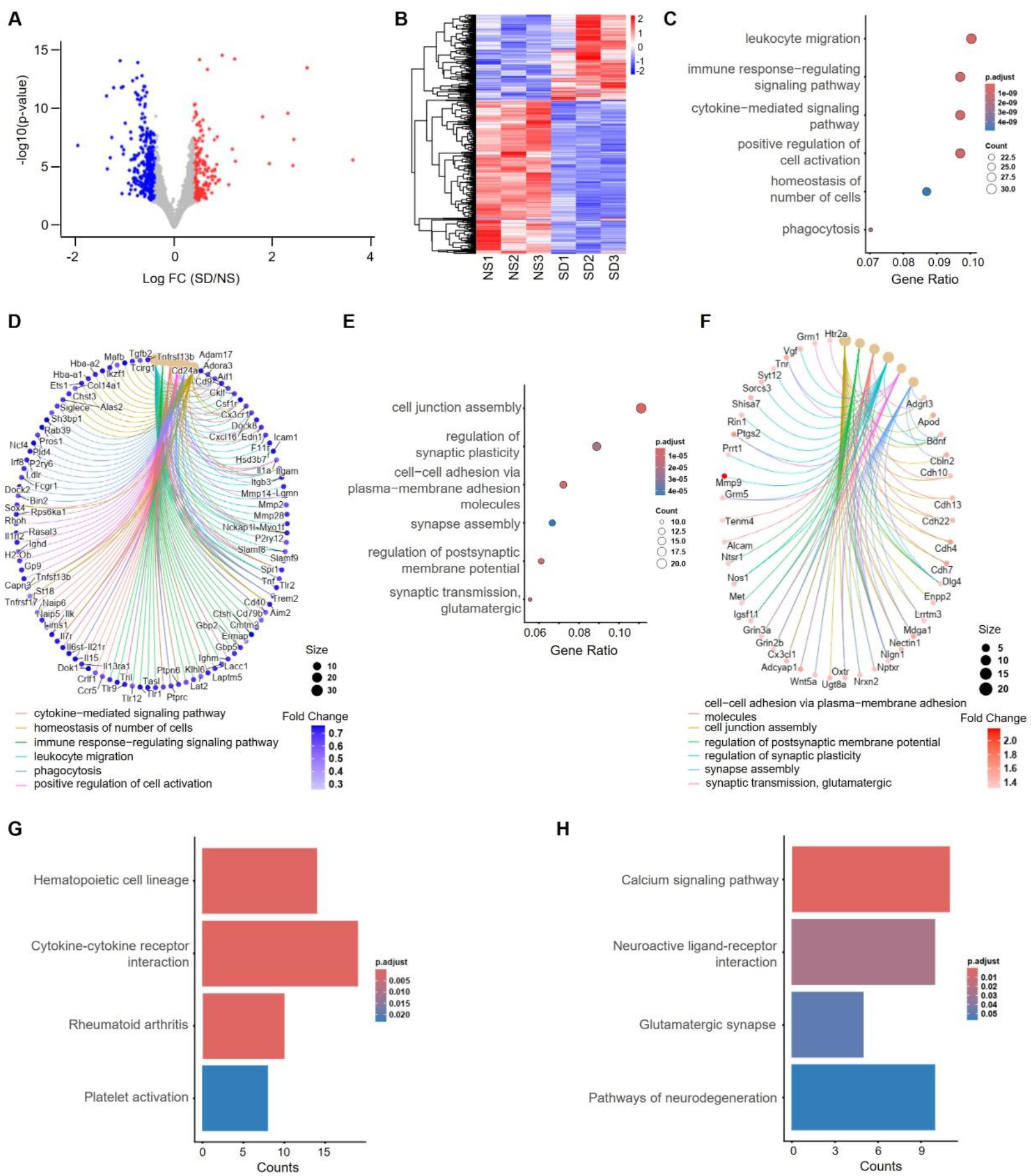
Transcriptional changes in neurons of cortex after sleep deprivation. (**A**) Volcano plot showing changes in gene expression for Sleep Deprived (SD)/Normal Sleep (NS) comparisons. Unpaired t-tests followed by false discovery rate (FDR) analysis were used to compare groups. Significantly downregulated (log2(SD/NS) < -0.4, FDR < 0.05) and upregulated (log2(SD/NS) > 0.4, FDR < 0.05) gene expression is color-coded with blue and red, respectively. Genes in grey are not significantly changed after sleep deprivation. (**B**) Heatmap showing differentially expressed genes between NS and SD conditions. (**C, E**) Dot plots showing significantly enriched top downregulated (C) and upregulated (E) biological processes. Circle size is proportional to gene count and the level of significance (Benjamini-Hochberg adjusted *P* value < 0.05) is color-coded. (**D, F**) Cnet plots show significantly enriched top-downregulated (D) and upregulated (F) biological processes and their gene linkages. Circle size is proportional to gene counts for biological processes. The edge is color-coded as a biological process. The fold change of downregulated and upregulated genes is color-coded with blue and red gradients, respectively. (**G, H**) Bar plots showing significantly enriched top downregulated (G) and upregulated (H) KEGG pathways. The level of significance (Benjamini-Hochberg adjusted *P* value < 0.05) is color-coded.

**FIGURE 3.**
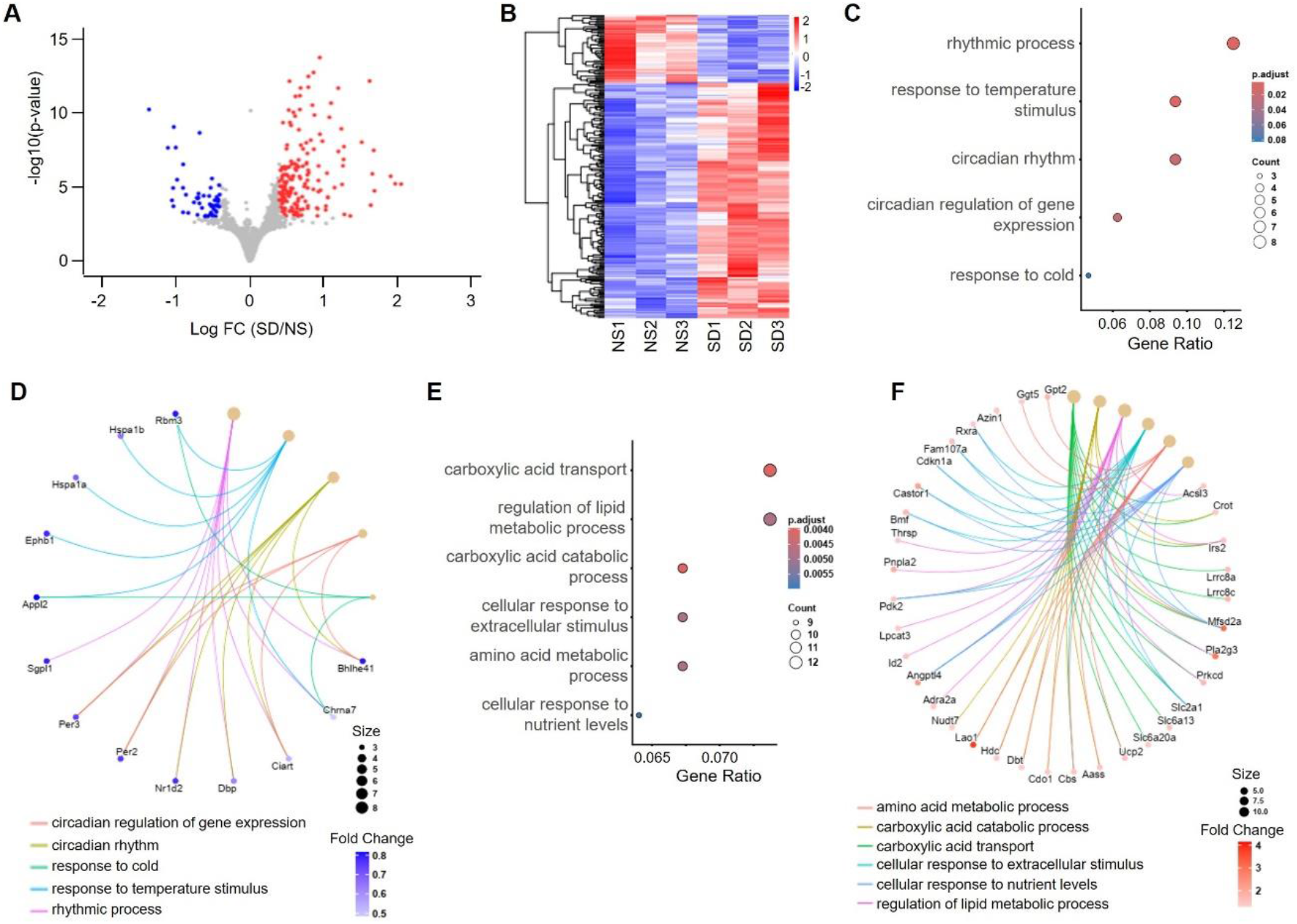
Transcriptional changes in astrocytes of cortex after sleep deprivation. (**A**) Volcano plot showing changes in gene expression for Sleep Deprived (SD)/Normal Sleep (NS) comparisons. Unpaired t-tests followed by false discovery rate (FDR) analysis were used to compare groups. Significantly downregulated (log2(SD/NS) < -0.4, FDR < 0.05) and upregulated (log2(SD/NS) > 0.4, FDR < 0.05) gene expression is color-coded with blue and red, respectively. Genes in grey are not significantly changed after sleep deprivation. (**B**) Heatmap showing differentially expressed genes between NS and SD conditions. (**C, E**) Dot plots showing significantly enriched top downregulated (C) and upregulated (E) biological processes. Circle size is proportional to gene count and the level of significance (Benjamini-Hochberg adjusted *P* value < 0.05) is color-coded. (**D, F**) Cnet plots show significantly enriched top-downregulated (D) and upregulated (F) biological processes and their gene linkages. Circle size is proportional to gene counts for biological processes. The edge is color-coded as a biological process. The fold change of downregulated and upregulated genes is color-coded with blue and red gradients, respectively.

**Table 1.**
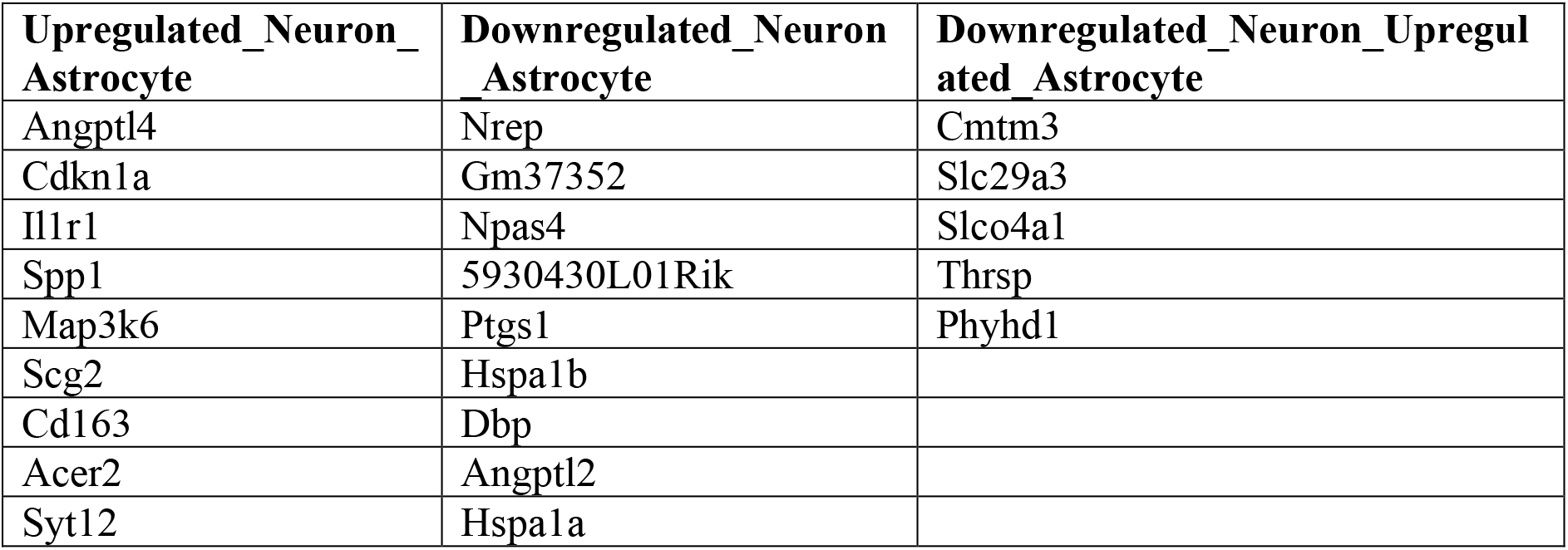
1.3-fold cutoff, FDR < 0.05.

| Upregulated_Neuron_Astrocyte | Downregulated_Neuron_Astrocyte | Downregulated_Neuron_Upregulated_Astrocyte |
| --- | --- | --- |
| Angptl4 | Nrep | Cmtm3 |
| Cdkn1a | Gm37352 | Slc29a3 |
| Il1r1 | Npas4 | Slco4a1 |
| Spp1 | 5930430L01Rik | Thrsp |
| Map3k6 | Ptgs1 | Phyhd1 |
| Scg2 | Hspa1b |  |
| Cd163 | Dbp |  |
| Acer2 | Angptl2 |  |
| Syt12 | Hspa1a |  |

Given these findings, we examined the biological processes affected by sleep deprivation within these cell types by performing gene-annotation enrichment analysis with the genes differentially downregulated and upregulated, separately. The downregulated genes were enriched for the leukocyte migration, immune response-regulating signaling pathway, cytokine−mediated signaling pathway and other inflammatory responses in neurons and rhythmic processes, circadian regulation of gene expression, and response to temperature in astrocytes (Fig. 2 C, D Fig. 3 C, D). On the other hand, upregulated genes were enriched for cell junction assembly, regulation of synaptic plasticity, post-synaptic membrane potential, and synapse assembly processes in neurons and carboxylic acid transport, catabolic process, and cellular response to extracellular stimulus in astrocytes (Fig. 2 E, F, Fig. 3 E, F). These findings indicate that sleep need differentially impacts cellular functions in neurons and astrocytes. We next performed KEEG pathway analysis and reported that downregulated genes were enriched for signaling of hematopoietic cell lineage, cytokine-cytokine receptor interaction, and other inflammatory pathways and upregulated ones were for calcium signaling, neuroactive ligand-receptor interaction, glutamatergic synapse and pathways of neurodegeneration in neurons (Fig. 2 G, H). This enrichment was not noticeable in astrocytes.

### Sleep deprivation alters nuclear protein abundance of neurons and astrocytes in the cortex

To evaluate how sleep deprivation influences the nuclear proteome of neurons and astrocytes, we performed quantitative proteomics of nuclei isolated from the cortical cell suspensions by using multiplex tandem mass tag (TMT) labeling coupled with liquid chromatography-tandem mass spectrometry (LC-MS/MS) (Fig. 1B; see Methods).

Sleep deprivation altered a higher proportion of nuclear proteins in neurons (70%, 734/1107) than in astrocytes (44%, 469/1063) (FDR < 0.05) (Fig. 4A-B, Fig. 5A-B). We previously reported similar alterations in total protein levels in neurons and astrocytes ^13^. Moreover, this trend is consistent with transcriptional changes in neurons and astrocytes we report here (see above). These analyses indicate that neurons are more sensitive to sleep deprivation than astrocytes. We applied 30%-fold-change (1.3-fold cutoff, FDR < 0.05) filtration for the magnitude of change in protein abundance and reported majority of proteins show higher abundance in the nuclei of neurons (319 upregulated and 278 downregulated) and lower abundance in the nuclei of astrocytes (128 upregulated and 181 downregulated). Next, we checked the directionality of the protein abundance in neurons and astrocytes. Interestingly, 191 proteins exhibited the same directionality in both cell types, while 5 showed opposite trends (Table 2). These results indicate that, unlike transcriptomic changes, the shifts in nuclear proteomes due to sleep loss are predominantly unidirectional.

**FIGURE 4.**
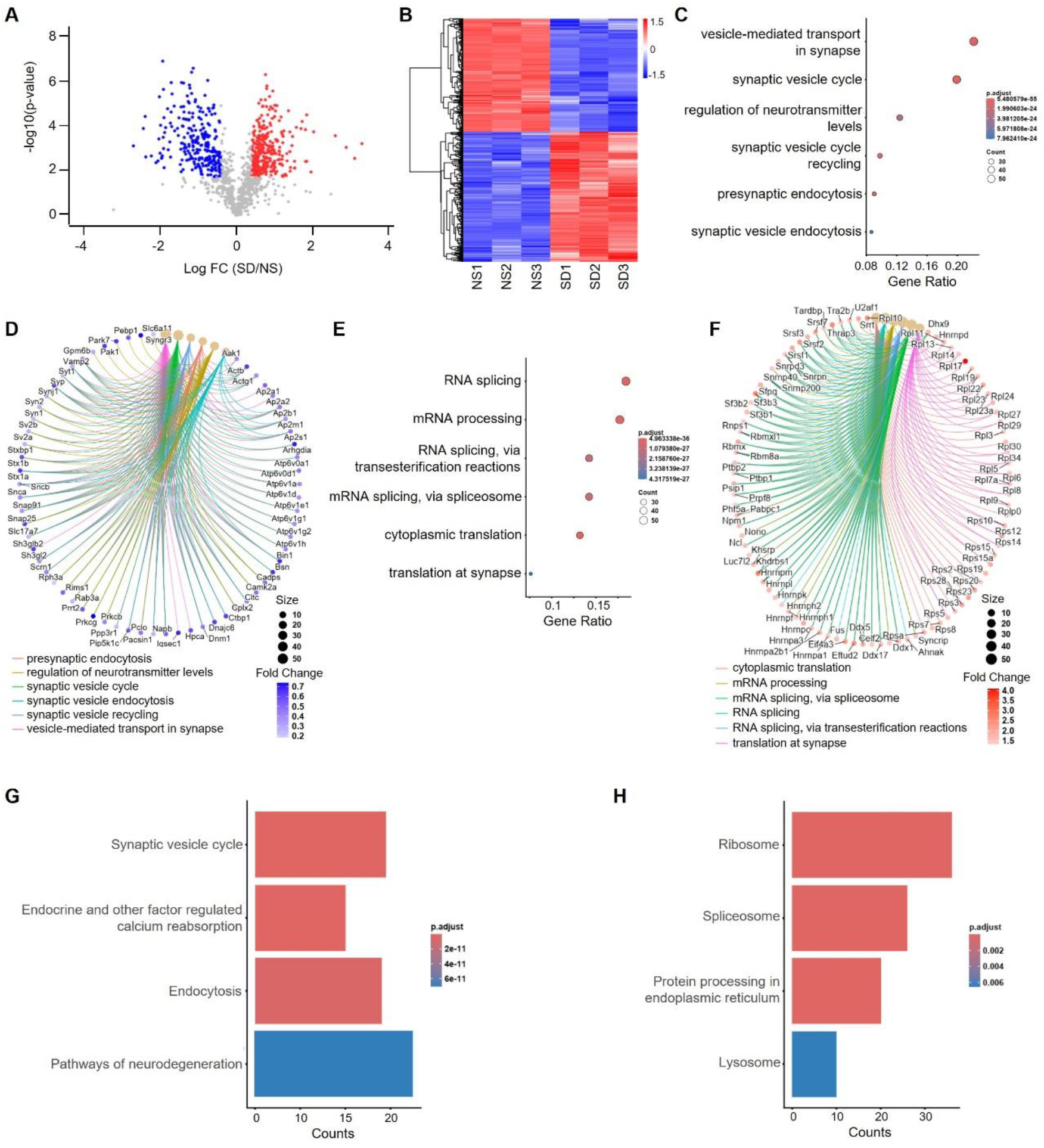
Sleep deprivation alters the abundance of nuclear proteins in neurons. (**A**) Volcano plot showing changes in protein expression for Sleep Deprived (SD)/Normal Sleep (NS) comparisons. Unpaired t-tests followed by false discovery rate (FDR) analysis were used to compare groups. Significantly downregulated (log2(SD/NS) < -0.4, FDR < 0.05) and upregulated (log2(SD/NS) > 0.4, FDR < 0.05) protein expression is color-coded with blue and red, respectively. Proteins in grey are not significantly changed after sleep deprivation. (**B**) Heatmap showing differentially expressed proteins between NS and SD conditions. (**C, E**) Dot plots showing significantly enriched top downregulated (C) and upregulated (E) biological processes. Circle size is proportional to protein count and the level of significance (Benjamini-Hochberg adjusted *P* value < 0.05) is color-coded. (**D, F**) Cnet plots show significantly enriched top-downregulated (D) and upregulated (F) biological processes and their protein linkages. Circle size is proportional to protein counts for biological processes. The edge is color-coded as a biological process. The fold change of downregulated and upregulated proteins is color-coded with blue and red gradients, respectively. (**G, H**) Bar plots showing significantly enriched top downregulated (G) and upregulated (H) KEGG pathways. The level of significance (Benjamini-Hochberg adjusted *P* value < 0.05) is color-coded.

**FIGURE 5.**
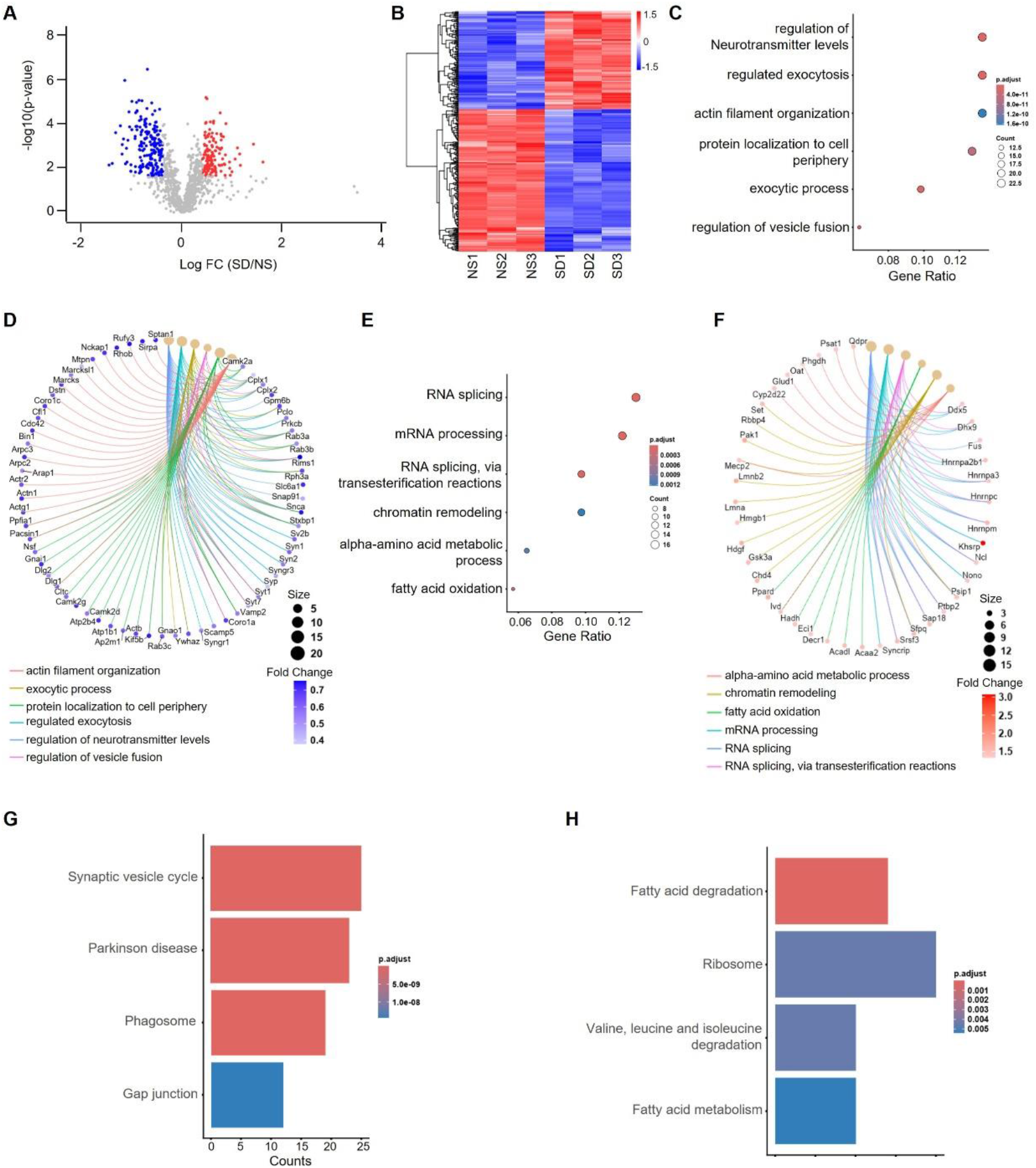
Sleep deprivation alters the abundance of nuclear proteins in astrocytes. (**A**) Volcano plot showing changes in protein expression for Sleep Deprived (SD)/Normal Sleep (NS) comparisons. Unpaired t-tests followed by false discovery rate (FDR) analysis were used to compare groups. Significantly downregulated (log2(SD/NS) < -0.4, FDR < 0.05) and upregulated (log2(SD/NS) > 0.4, FDR < 0.05) protein expression is color-coded with blue and red, respectively. Proteins in grey are not significantly changed after sleep deprivation. (**B**) Heatmap showing differentially expressed proteins between NS and SD conditions. (**C, E**) Dot plots showing significantly enriched top downregulated (C) and upregulated (E) biological processes. Circle size is proportional to protein count and the level of significance (Benjamini-Hochberg adjusted *P* value < 0.05) is color-coded. (**D, F**) Cnet plots show significantly enriched top-downregulated (D) and upregulated (F) biological processes and their protein linkages. Circle size is proportional to protein counts for biological processes. The edge is color-coded as a biological process. The fold change of downregulated and upregulated proteins is color-coded with blue and red gradients, respectively. (**G, H**) Bar plots showing significantly enriched top downregulated (G) and upregulated (H) KEGG pathways. The level of significance (Benjamini-Hochberg adjusted *P* value < 0.05) is color-coded.

**Table 2.**
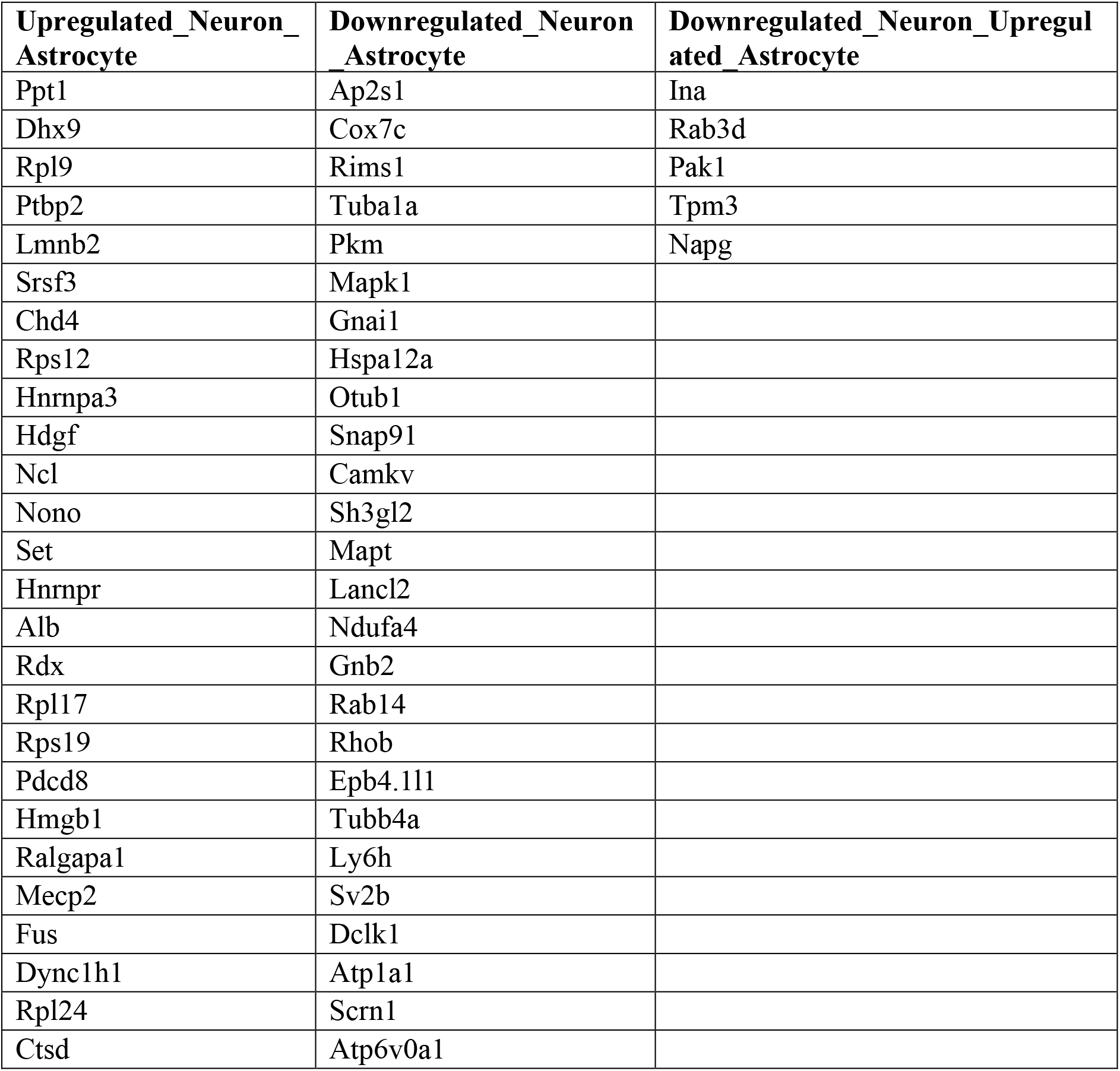
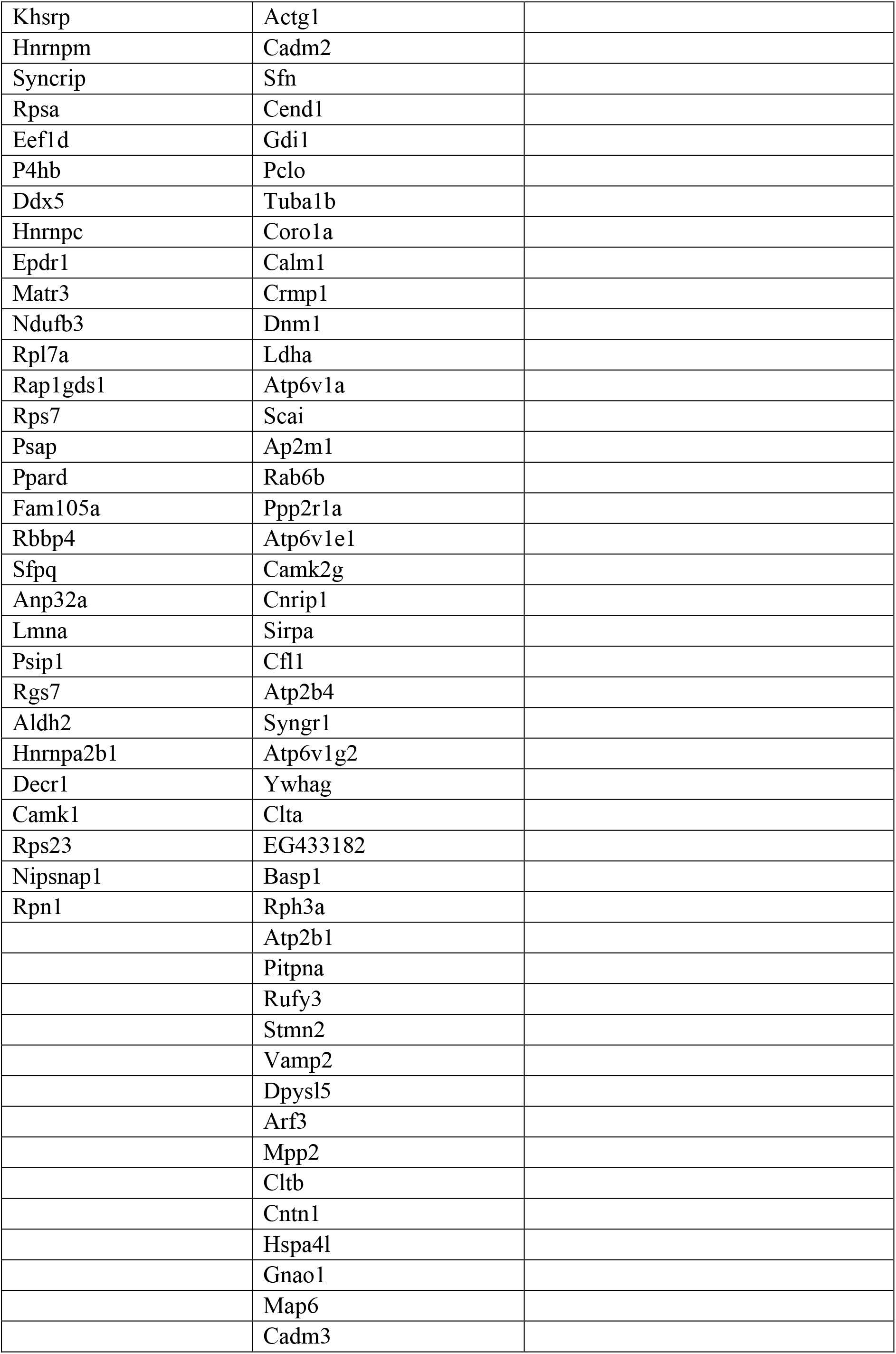
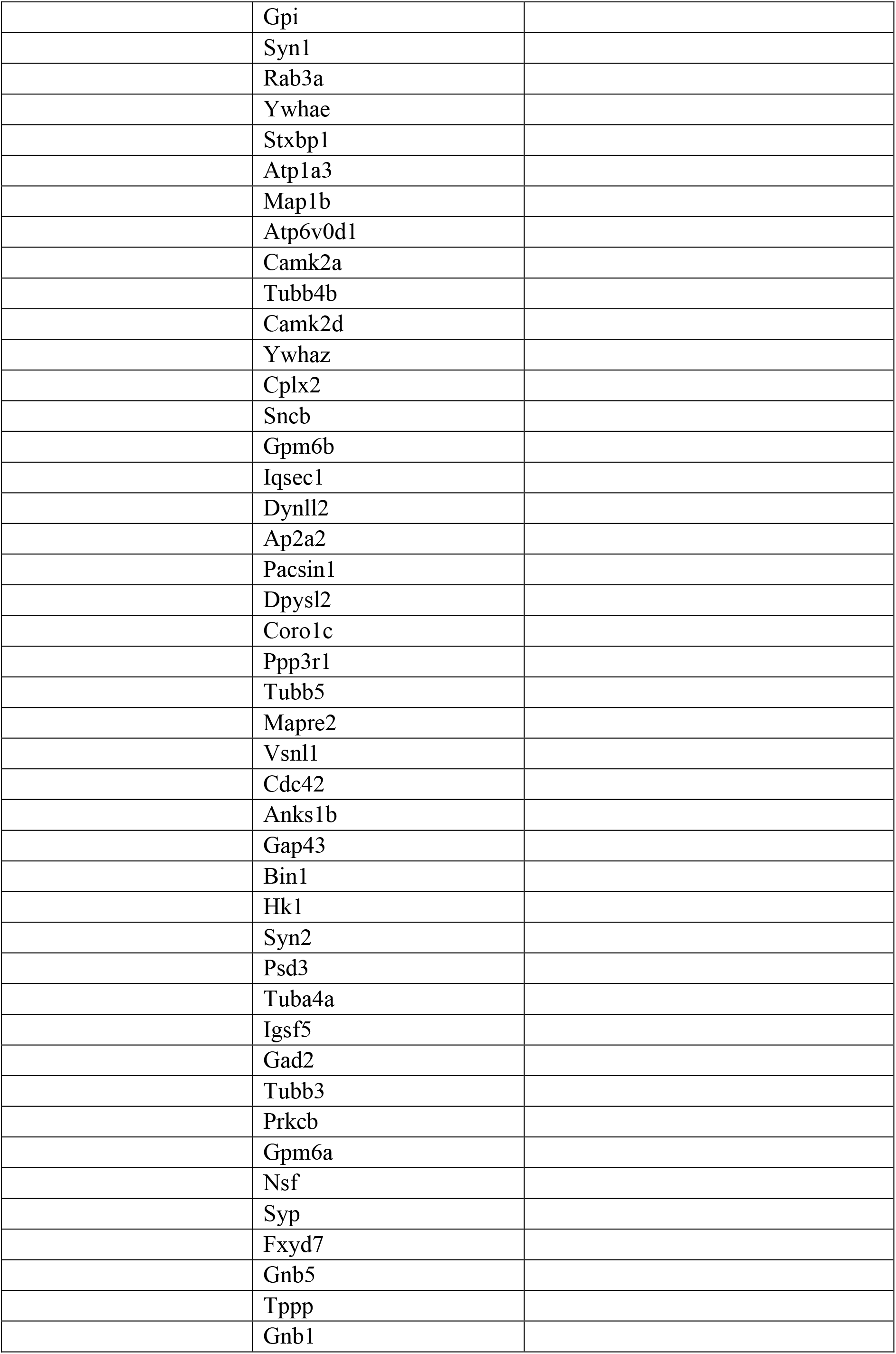
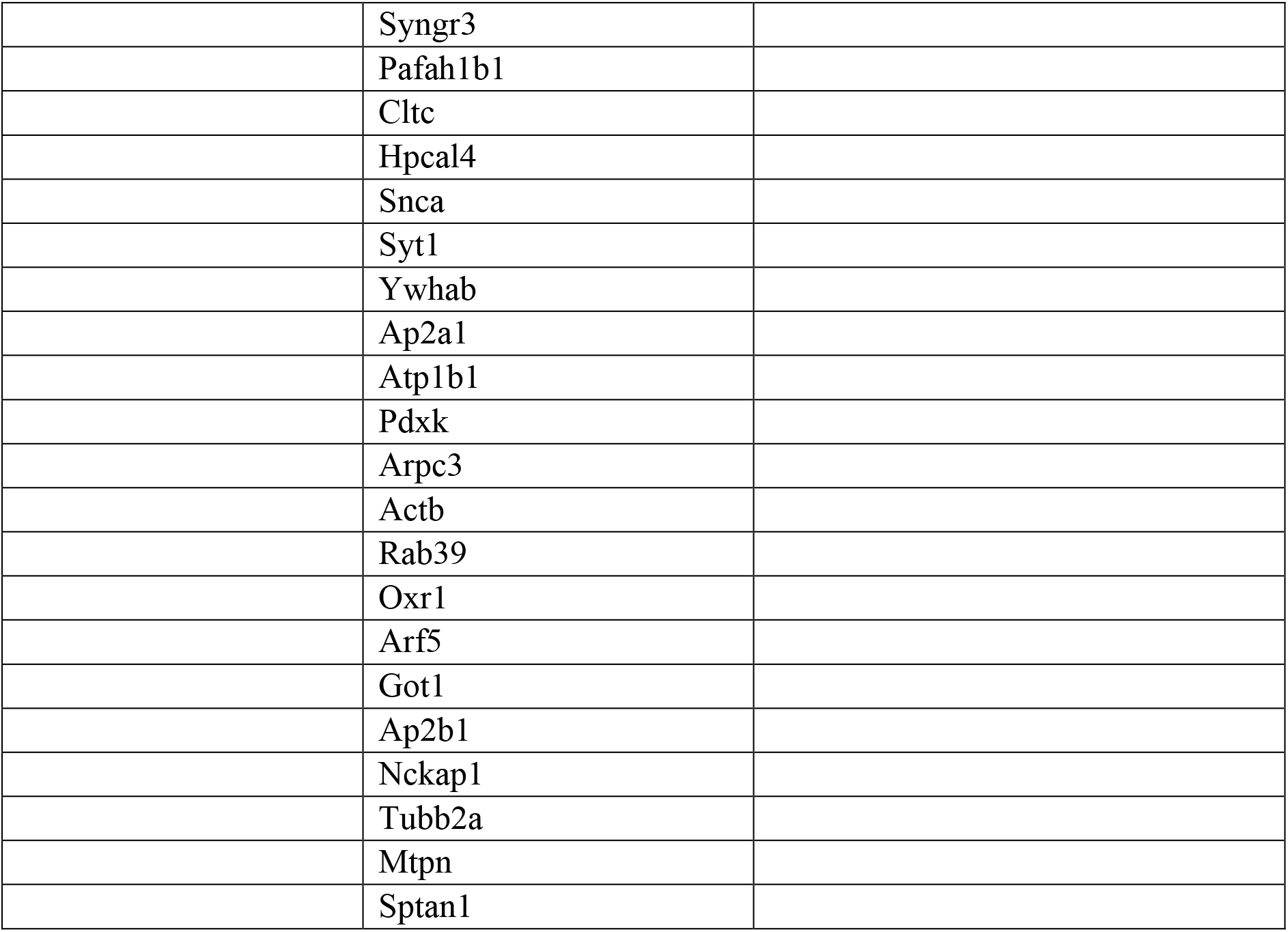
1.3-fold cutoff, FDR < 0.05.

To understand the biological functions of these alterations, we performed gene-annotation enrichment analysis separately on proteins downregulated and upregulated. These analyses revealed that downregulated proteins were enriched for vesicle-mediated transport at the synapse, regulation of neurotransmitter levels, synaptic vesicle and pre-synaptic endocytosis, and synaptic vesicle cycle recycling in neurons (Fig. 4 C, D). In astrocytes, enriched terms pointed to the regulation of exocytosis, actin filament organization, protein localization to the cell periphery, and regulation of neurotransmitter levels and vesicle fusions (Fig. 5 C, D). Upregulated proteins were enriched for RNA metabolism in both cell types, with cytoplasmic translation enriched in neurons and carboxylic acid catabolism and amino acid metabolic process in astrocytes (Fig. 4E, F, Fig. 5E, F). Overall, these analyses indicate that sleep deprivation similarly affects nuclear protein expression related to RNA metabolism in both cell types whereas proteins related to organic acid metabolism, amino acid metabolism, neurotransmitter transport, and vesicle-mediated transport are expressed differently within the nuclei of neurons and astrocytes. The KEGG pathways analysis reveals that downregulated proteins enriched for signaling of synaptic vesicle cycle, calcium reabsorption, endocytosis, and other pathways of neurodegeneration and upregulated ones were for the ribosome, spliceosome, protein processing in the endoplasmic reticulum and lysosomes in neurons (Fig. 4 G, H). In astrocytes, these enrichments were for the synaptic vesicle cycle, Parkinson’s disease, phagosome, and gap junction with the downregulated proteins whereas upregulated proteins enriched for the pathways associated with fatty acid and branch chain amino acid (BCAA) (Valine, leucine, and isoleucine) degradation (Fig. 5 G, H).

### Identification of transcription factor (TF) binding site motifs in SD-responsive genes

Next, we asked how sleep loss might regulate gene expression changes in neurons and astrocytes. To answer this question, we performed gene regulatory region analysis of 1000 bp of DNA proximal to the transcription start site (TSS) ^22^. We identified over-represented TF binding site motifs in significantly altered genes (1.3-fold cutoff, FDR < 0.05). In neurons, 30 and 40 TFs were enriched among downregulated and upregulated genes, respectively. In astrocytes, 22 and 89 TFs were enriched among downregulated and upregulated genes, respectively (FDR (Benjamini) < 0.05) (Table 3).

**Table 3.**
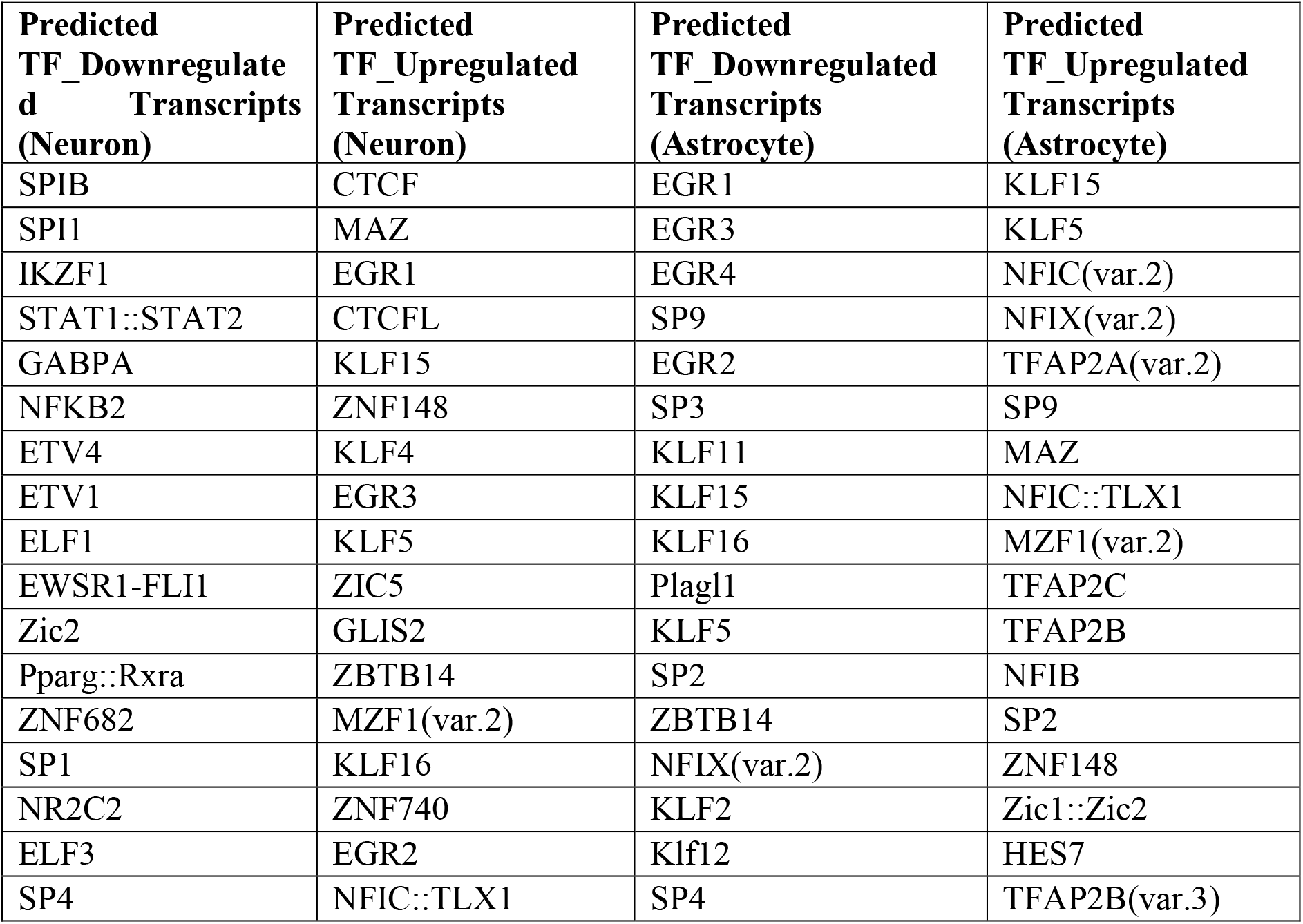
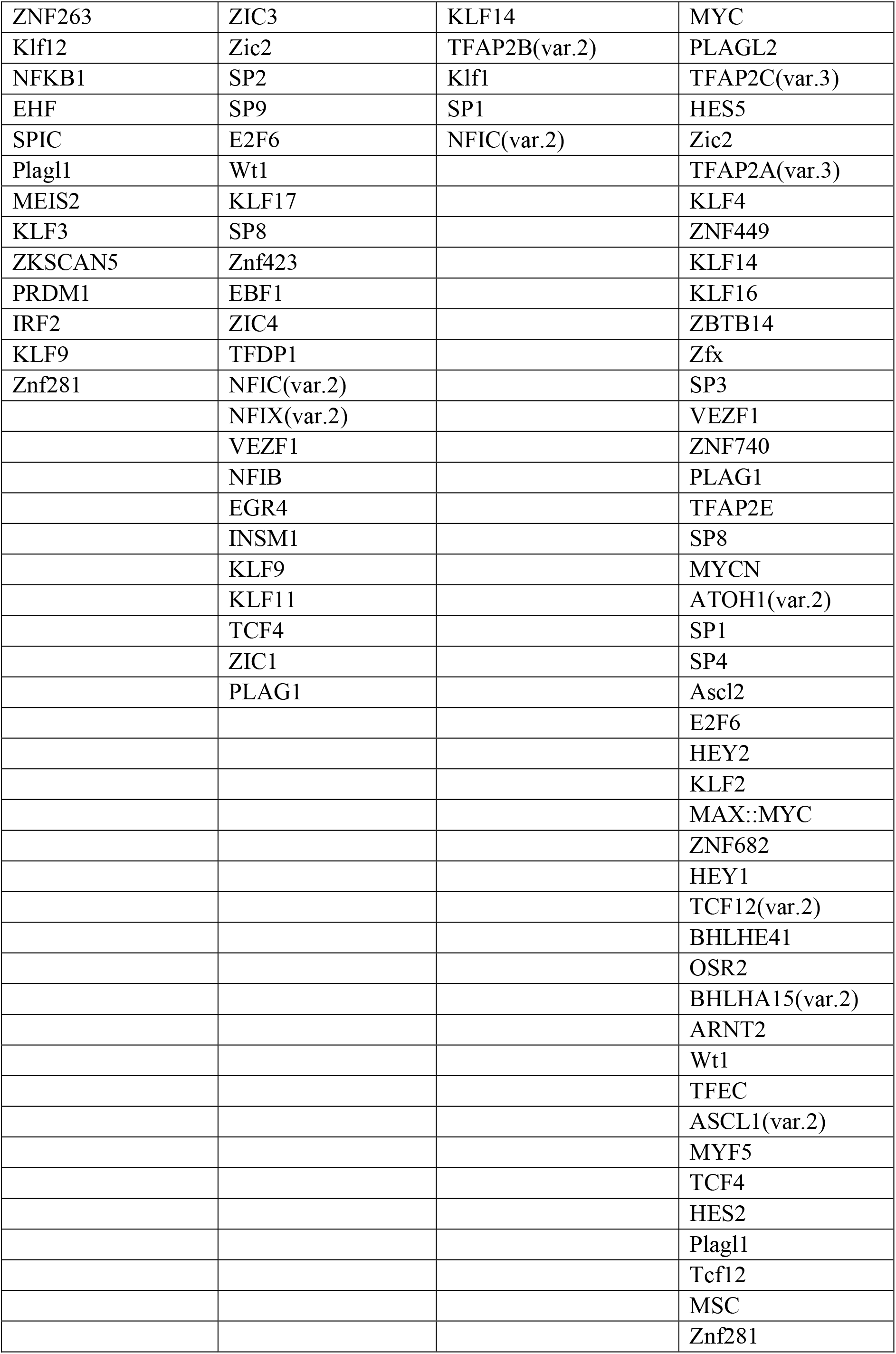
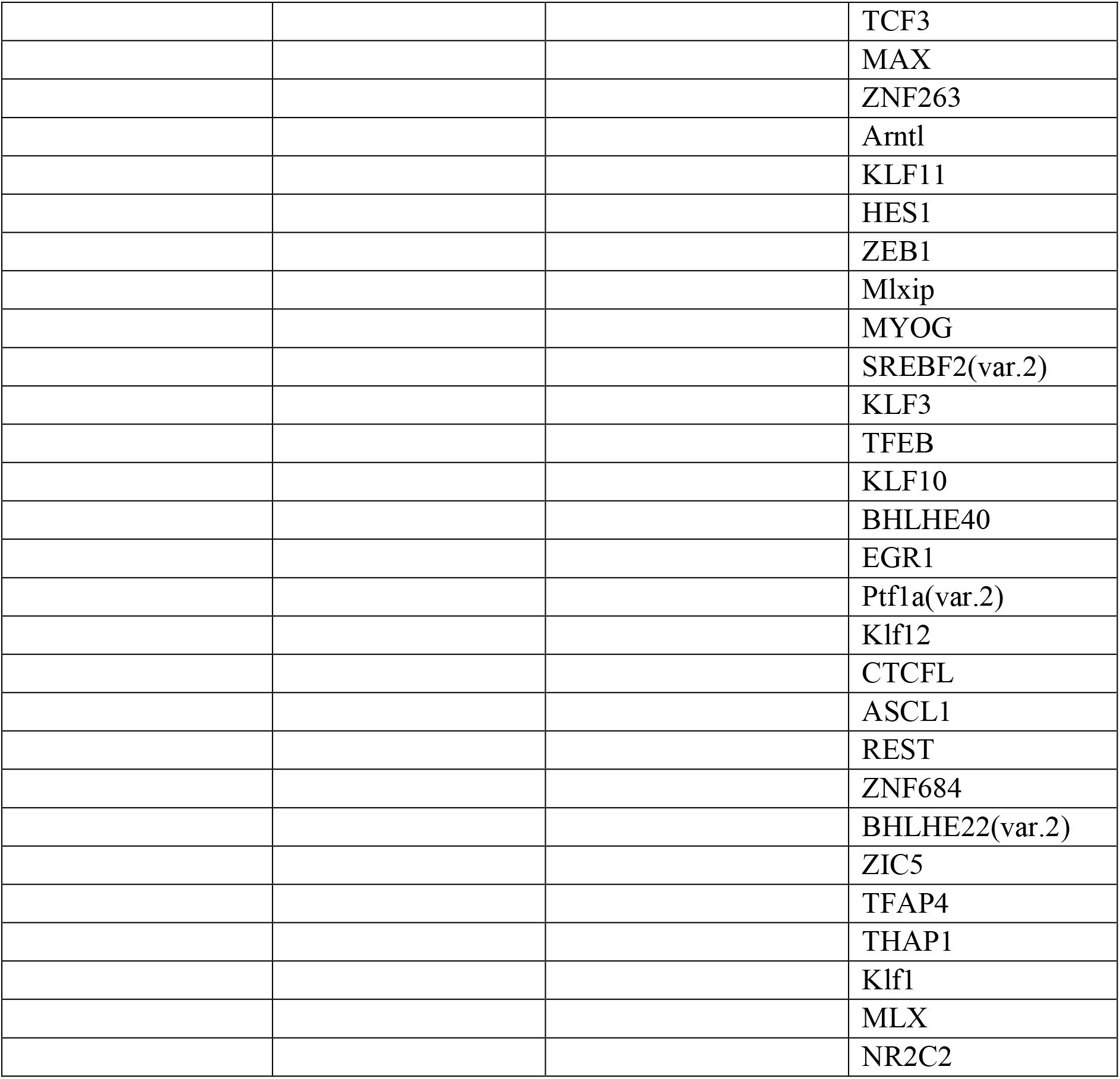
FDR < 0.05.

Top-scoring TFs enriched among downregulated neuronal genes, such as *Spib* (Transcription factor Spi-B), *Spi1* (Transcription factor PU.1), and *Stat1* (*Signal transducer and activator of transcription 1*), play roles in immune and inflammatory responses (Fig. 6A) ^23,24^. TFs enriched among upregulated neuronal genes included *Ctcf* (Transcriptional repressor CTCF), *Maz* (Myc-associated zinc finger protein), and *Egr1* (Early growth response protein 1), with *Klf15* (Krueppel-like factor 15) and *Egr1* previously reported in sleep and circadian rhythm studies (Fig. 6B) ^25,26^.

**FIGURE 6.**
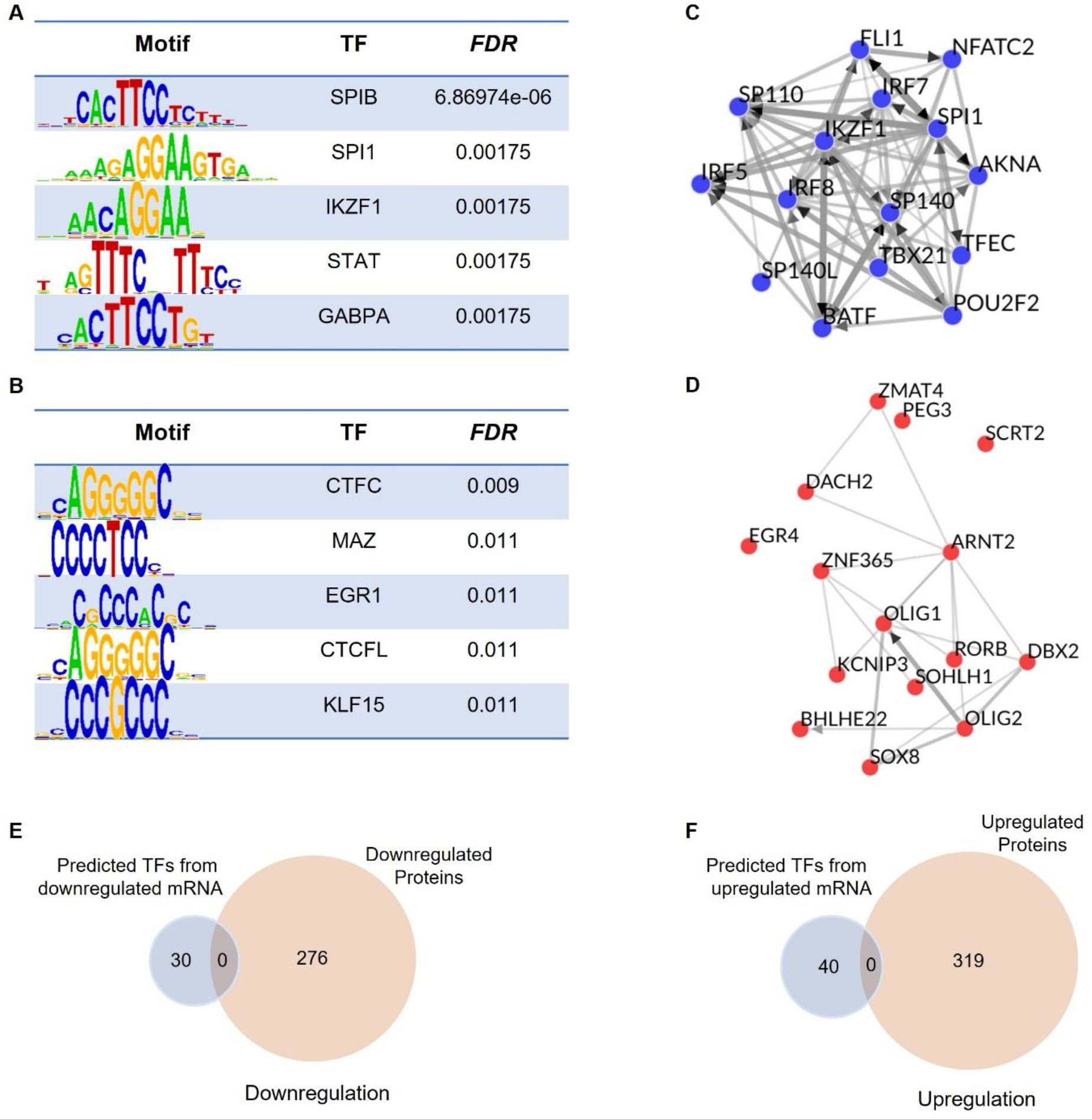
Transcription factor (TF) binding site motifs of the sleep-responsive genes in neurons. (**A-B**) Top sequence motif (FDR < 0.01) of the transcriptional regulators for the sleep-responsive significantly (FDR < 0.05) downregulated (log2(SD/NS) < -0.4) (A) and upregulated (log2(SD/NS) > 0.4) (B) transcripts. (**C-D**) Transcription factor co-regulatory networks of sleep-responsive significantly (FDR < 0.05) downregulated (log2(SD/NS) < -0.4) (C) and upregulated (log2(SD/NS) > 0.4) (D) transcripts. (**E-F**) Venn diagram showing the overlap between predicted downregulated (E) and upregulated (F) TFs and proteins in neurons. NS: Normal Sleep, SD: Sleep Deprived

In astrocytes, *Sp9* (Transcription factor Sp9) and EGR family TFs (*Egr1, Egr2, Egr3, Egr4*) were enriched among downregulated genes (Fig. 7A), while *Klf15*, *Nfic* (Nuclear factor 1 C-type), and *Tfap2A* (Transcription factor AP-2-alpha) were enriched among upregulated genes (Fig. 7B). *Tfap2A* has been implicated in NREM sleep regulation ^27^.

**FIGURE 7.**
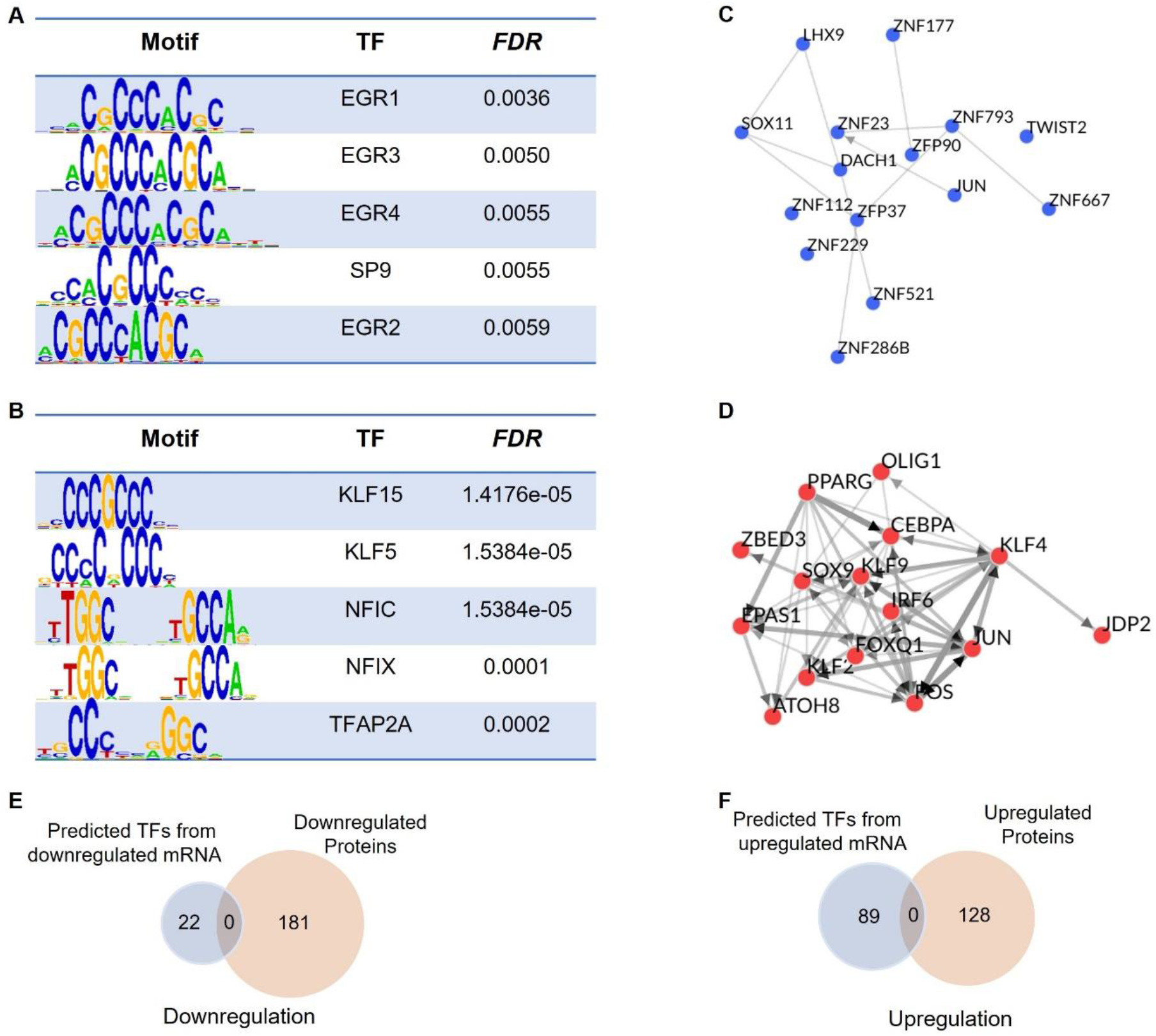
Transcription factor (TF) binding site motifs of the sleep-responsive genes in astrocytes. (**A-B**) Top sequence motif (FDR < 0.01) of the transcriptional regulators for the sleep-responsive significantly (FDR < 0.05) downregulated (log2(SD/NS) < -0.4) (A) and upregulated (log2(SD/NS) > 0.4) (B) transcripts. (**C-D**) Transcription factor co-regulatory networks of sleep-responsive significantly (FDR < 0.05) downregulated (log2(SD/NS) < -0.4) (C) and upregulated (log2(SD/NS) > 0.4) (D) transcripts. (**E-F**) Venn diagram showing the overlap between predicted downregulated (E) and upregulated (F) TFs and proteins in astrocytes. NS: Normal Sleep, SD: Sleep Deprived

We analyzed TF-TF co-regulatory networks by using ChIP-X Enrichment Analysis 3 (ChEA3) ^28^ (Fig. 6 C, D; Fig. 7 C, D). Downregulated TFs in neurons showed stronger interactions than upregulated ones (Fig. 6 C, D), whereas in astrocytes this interaction was robust in upregulated TF (Fig. 7 C, D). This analysis allowed us to understand the other TFs that share targets with enriched TFs. Next, we searched for these predicted TFs in the nuclear proteomics data set by performing an overlap between the predicted downregulated and upregulated TFs with proteins that similarly altered in both these cell types (Fig. 6 E, F; Fig. 7 E, F). However, predicted TFs were not found in the nuclear proteomics data set (Fig. 6E, F; Fig. 7E, F), suggesting a discrepancy between gene expression and functional protein abundance ^16,17^. Moreover, since the transcriptomes and proteomes were sampled at the same time-point, mRNA inductions may not be translated into protein abundance changes immediately.

### Characterization of SD-responsive TFs in nuclear proteomes of neurons and astrocytes

To gain insights into the regulatory mechanisms underlying the transcriptional changes observed in sleep-deprived neurons and astrocytes, we characterized significantly altered nuclear proteins (1.3-fold cutoff, FDR < 0.05) using the TRRUST database ^29^ and identified TFs in both cell types. In neurons, 27 TFs were upregulated and 2 downregulated (Fig. 8A), while in astrocytes, 8 TFs were upregulated and 2 downregulated (Fig. 8B). The higher number of altered TFs in neurons suggests that they may be more sensitive to sleep deprivation-induced transcriptional changes compared to astrocytes.

**FIGURE 8.**
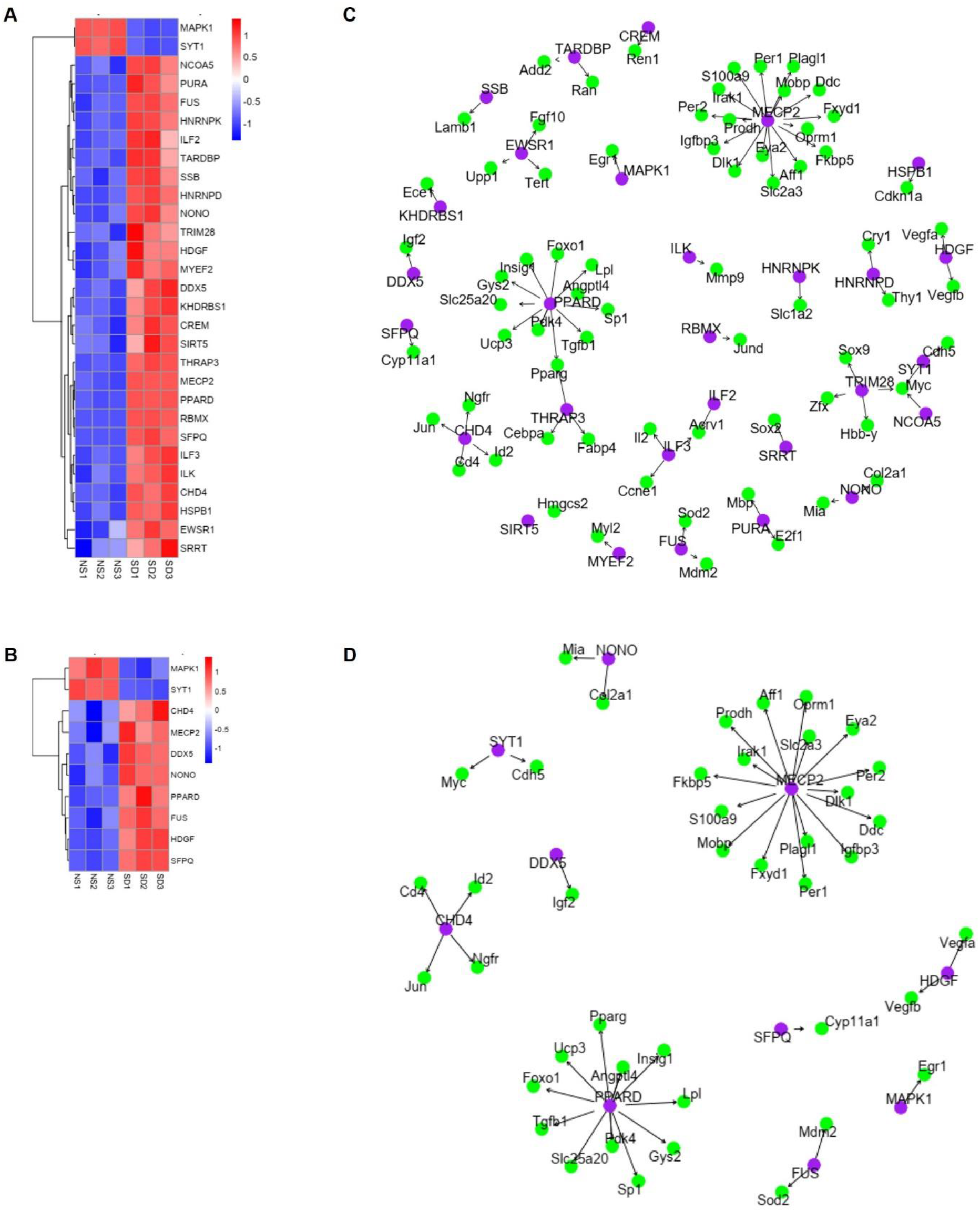
Sleep deprivation alters the abundance of nuclear transcription factors (TFs) in neurons and astrocytes. **(A-B)** Heatmap showing differentially expressed TFs between NS and SD conditions in neurons (A) and astrocytes (B). Unpaired t-tests followed by false discovery rate (FDR) analysis were used to compare groups. Significantly downregulated (log2(SD/NS) < -0.4, FDR < 0.05) and upregulated (log2(SD/NS) > 0.4, FDR < 0.05) TF expression is color-coded with blue and red, respectively. (**C-D**) Network plot of nuclear TF and their co-expressed target genes for neurons (C) and astrocytes (D). Purple and green circles represent TF and target genes, respectively. NS: Normal Sleep, SD: Sleep Deprived

TRRUST network analysis revealed that in neurons, MECP2 regulates the expression of *Mobp*, *Igfbp3*, and *S100a9* (Fig. 8C). MECP2 is a crucial regulator of neuronal function and has been implicated in neurodevelopmental disorders such as Rett syndrome ^30^. Its role in sleep deprivation-induced gene expression changes highlights the potential impact of sleep loss on neuronal function. PPARD, another TF altered in neuronal nuclei, regulates *Angptl4* and *Pdk4*, which are involved in lipid metabolism and glucose homeostasis, respectively ^31,32^. This suggests that sleep deprivation may influence energy metabolism in neurons through PPARD-mediated transcriptional regulation. Additionally, ILK and SIRT5 regulate genes involved in extracellular matrix remodeling (*Mmp9*) and ketogenesis (*Hmgcs2*), respectively ^33,34^. These findings indicate that sleep deprivation may affect a wide range of cellular processes in neurons through the action of specific TFs.

In astrocytes, PPARD and CHD4 regulate *Angptl4* and *Id2*, respectively (Fig. 8D). The presence of PPARD in both neuronal and astrocytic nuclei suggests that it may play a role in the sleep deprivation response across different cell types. CHD4, a chromatin remodeling factor, regulates *Id2*, which is involved in cell cycle progression and differentiation ^35^. This indicates that sleep deprivation may influence astrocyte proliferation and differentiation through CHD4-mediated transcriptional control.

## Discussion

This study provides the first comprehensive analysis of gene expression and nuclear protein abundance in neurons and astrocytes of the cortex during sleep loss. Our findings reveal distinct responses to sleep deprivation in these two key cell types involved in sleep regulation. Through integrated analysis of gene regulatory processes and transcriptional and translational shifts, we identified cell type-specific transcription factors (TFs) that regulate gene expression programs differently during sleep loss challenges. These results highlight key TFs and their target genes as important candidates for exploring the molecular mechanisms of sleep regulation in a cell type-specific manner.

Neurons and astrocytes in the mammalian cortex are integral to the neural mechanisms driving homeostatic sleep propensity ^9,36–38^. Our combined transcriptomics and nuclear proteomics approach provides novel insights into sleep drive correlates within these two cell populations. Transcriptome analysis revealed that neurons responded to sleep deprivation at larger proportions than astrocytes, consistent with our previous findings ^13^. This suggests that cortical neurons are the primary responders to sleep loss and homeostatic sleep need, while astrocytes play a critical but secondary role in maintaining sleep homeostasis.

Interestingly, we observed directional differences in gene expression changes between neurons and astrocytes. Under high sleep need, neurons generally decreased expression of most altered genes, while astrocytes increased expression. The lack of consistent changes between these cell types underscores the importance of single-cell resolution analysis for understanding sleep-related transcriptional patterns.

Functional analysis revealed that sleep loss reduces biological processes related to immune and inflammatory responses in neurons and circadian rhythm maintenance in astrocytes. The attenuation of immune and inflammatory responses in neurons aligns with the known rapid antidepressant effects of acute sleep deprivation ^39,40^. Neurons employ various strategies to reduce immune and inflammatory responses, including the release of anti-inflammatory neurotransmitters and brain-derived neurotrophic factor (BDNF), and the inhibition of inflammatory signaling ^41–43^. These mechanisms may represent a protective response to mitigate inflammatory damage following prolonged wakefulness.

The reduction in circadian rhythm-related gene expression in astrocytes suggests a potential mechanism for the feedback effect of sleep deprivation on the circadian pacemaker ^44,45^. This finding is supported by previous studies showing that sleep deprivation modulates clock gene expression in the cortex and that cortical astrocytes regulate the sleep-wake cycle ^46,47^.

Conversely, sleep deprivation increased synapse assembly and synaptic transmission-related pathways in neurons and organic acid catabolism in astrocytes. The neuronal changes align with the synaptic homeostasis hypothesis of sleep-wake regulation, which posits that sleep functions to downscale or renormalize synapses ^48^. The astrocytic changes suggest a role in facilitating the rebuilding and synthesis of macromolecules used during waking hours ^49^, highlighting the compartmentalization of sleep functions between these cell types.

Our analysis of transcriptomics and nuclear proteomics datasets revealed cell type-specific TFs mediating gene expression changes during forced wakefulness. In neurons, we observed downregulation of SPIB, SPI1, IKZF1, STAT1, GABPA, SYT1, and MAPK, and upregulation of CTCF, MAZ, EGR1, CTCFL, KLF15, MECP2, PPARD, HSPB1, ILK, and SIRT5. In astrocytes, SP9, EGRs, MAPK1, and SYT1 were downregulated, while KLF15, KLF5, NFIC, NFIX, and TFAP2A were upregulated. Several of these TFs, including SYT1, EGR1, MECP2, TFAP2A, and PPARD, have previously been implicated in sleep regulation and related pathologies ^27,50–54^. Our findings provide a comprehensive categorization of sleep loss-responsive TFs in neurons and astrocytes, laying the groundwork for future targeted investigations into their roles in sleep regulation.

The nuclear proteome analysis revealed consistent changes in RNA metabolism (RNA splicing and mRNA processing) in both cell types under higher sleep need. This suggests that disruption of RNA processing may be a key aspect of nuclear homeostatic maintenance during sleep loss^55^. The identification of RNA-binding proteins among the commonly characterized TFs (DDX5, FUS, NONO, SFPQ) further supports this notion ^56–59^. Contrary to RNA metabolism, accumulated sleep drive activates pathways related to proteostasis in neurons and the degradation of fatty acids and branched-chain amino acids (BCAAs) in astrocytes. Of note, the assumptions that sleep restores the intracellular order disrupted during sleep loss by protein homeostasis and synthesis of macromolecules are segregated as neuronal and astrocytic functions from our study ^49,60–62^. Additionally, sleep deprivation reduced vesicle-mediated transport in synapses and regulation of neurotransmitter levels in neurons, while affecting actin filament organization and protein localization to the cell periphery in astrocytes. These functional changes reinforce the crucial role of synaptic activity regulation in maintaining sleep-wake balance ^1,63^.

Thus, our multi-omics approach and analysis of gene regulatory networks demonstrate that neurons and astrocytes in the cortex respond differentially to accumulated sleep pressure through distinct gene regulatory processes and expression programs. These findings build upon our previous work showing that molecular determinants of sleep and wake vary within individual brain cell types ^13^. The comprehensive dataset generated in this study provides a valuable resource for further exploration of cell type-specific functions in sleep homeostasis.

While our study offers novel insights into the molecular mechanisms of sleep regulation, some limitations should be acknowledged. The use of bulk RNA sequencing and nuclear proteomics may obscure more subtle cell type-specific changes or rare cell populations. Future studies employing single-cell technologies could provide higher resolution data to address this limitation. Additionally, our focus on the cortex leaves open the question of how other brain regions respond to sleep loss. Comparative studies across multiple brain areas could yield a more comprehensive understanding of sleep regulation at the systems level.

Future directions for this work include targeted manipulation of the identified TFs to elucidate their causal roles in sleep regulation. Chromatin immunoprecipitation sequencing (ChIP-seq) or Cleavage Under Targets and Release Using Nuclease (CUT&RUN) sequencing experiments could further validate the predicted TF-target gene interactions. Moreover, investigating the temporal dynamics of these molecular changes across the sleep-wake cycle could provide insights into the kinetics of sleep homeostasis at the cellular level.

## Methods

### Animals

All animal studies adhered to approved University of Pennsylvania and ARRIVE guidelines. All animal experimental protocols were approved by the Institutional Animal Care and Use Committee (IACUC) at the Perelman School of Medicine at the University of Pennsylvania. Wild-type, male C57BL/6 J mice were procured from Jackson Laboratories and allowed to acclimate for at least two weeks before experiments. For sleep deprivation experiments, mice were housed individually in automated sleep fragmentation chambers (Model #80391, Campden/Lafayette Instrument Lafayette, IN, USA). At all times, the mice were given *ad libitum* access to food and water under standard housing conditions, under a 12-h light: 12-h dark cycle.

### Experimental design and sleep deprivation

Animals were divided into *ad libitum* Normal Sleep (NS) and Sleep Deprived (SD) groups. The NS group was left undisturbed SD group was sleep deprived by using a device that applied tactile stimulus with a horizontal bar sweeping just above the cage floor (bedding), as described previously ^13^. Once the sweeper was on, animals needed to step over it to continue their normal activities. Sleep deprivation began at the start of the light cycle (Zeitgeber Time 0 (ZT0)) and lasted for 12 h, with continuous sweeping mode (approximately 7.5 s cycle time). We made additional attempts to maintain wakefulness during the second half of the sleep deprivation period by occasionally tapping on the cage or gently touching the animals with a brush. Following sacrifice by cervical dislocation, whole brains were isolated at ZT12 (i.e. lights off). Isolated whole brains were placed in ice-cold Hibernate EB (BrainBits LLC, HEB) media, and the cerebral cortex was quickly dissected for the preparation of single-cell suspensions.

### Astrocyte and neuron separation from cortical single-cell suspension

To perform cell-type-specific nuclear transcriptomics and proteomics, astrocytes and neurons were separated from cortical single-cell suspensions as previously described ^64^ with slight modifications. Single-cell suspensions were incubated with FcR Blocking Reagent (Miltenyi Biotec, Cat # 130-092-575) and Anti-ACSA-2 MicroBeads (Miltenyi Biotec, Cat # 130-097-678) for 10-15 min at 2-8°C. Cells were spun down for 10 min. at 300 x g at room temp. and resuspended in a minimal volume of PBS with 0.5% of BSA before passing through an LS Column. Flow-through was collected for further neuron separation, the LS Column that retained astrocytes was removed from the magnetic field and the ACSA-2 labeled astrocytes eluted. Cells were pelleted from flow-through at 300 x g, 8 min, the supernatant carefully removed, and resuspended in a minimal volume of PBS with 0.5% of BSA. Suspensions were incubated with Non-Neuronal Cell Biotin-Ab cocktail (Miltenyi Biotec, Cat # 130-115-389) for 5 min and Anti-Biotin MicroBeads (Miltenyi Biotec, Cat # 130-090-485) for 10 min at 2-8°C, respectively. Following incubation, suspensions were passed through an LS Column placed in a magnetic field and non-labeled neuronal cells were collected in flow-through.

### Library preparation and RNA-sequencing

Astrocytes and neuron samples were lysed using TRIzol®, and RNA was extracted with the Direct-zol™ RNA MiniPrep Kit (Zymo Research), following the manufacturer’s instructions. 1μg of RNA was used to prepare total RNA libraries with the KAPA-RiboErase Hyper Prep kit, as per the manufacturer’s protocol. The libraries were sequenced on an Illumina HiSeq 4000, generating 100bp pair-ended reads. The reads were aligned to the Mouse genome GRCm39 using STAR version 2.7.10a. Transcript abundance was quantified as FPKM, TPM, and counts using StringTie v2.2.1. Genes with counts below 10 in all the samples were excluded from further analysis. Differential expression analysis was performed on the above filtered counts table using R package DESeq2 v1.44.0 to compare SD versus NS conditions. Log fold changes were adjusted using the *lfcShrink* function within the DESeq2 package. Genes with an FDR-adjusted p-value less than 0.05 were considered differentially expressed.

### TMT-based nuclear proteomics

Specific enrichment of nuclear proteins from astrocytes and neurons (NS, SD, 3 biological replicates) was performed using a Qproteome Nuclear Protein Kit (QIAGEN, Cat # 37582) following the manufacturer’s instructions. Subsequently, protein concentration in each sample was determined by using the Pierce™ BCA Protein Assay Kit (Thermo Fisher Scientific, Cat # 23225). 10-25µg protein per condition was transferred into new microcentrifuge tubes and 5 µL of the 200 mM TCEP was added to reduce the cysteine residues and the samples were then incubated at 55°C for 1-h. Subsequently, the reduced proteins were alkylated with 375 mM iodoacetamide (freshly prepared in 100 mM TEAB) for 30 min in the dark at room temperature. Then, trypsin (Trypsin Gold, Mass Spectrometry Grade; Promega, Cat # V5280) was added at a 1:40 (trypsin: protein) ratio and samples were incubated at 37°C for 12-h for proteolytic digestion. After in-solution digestion, peptide samples were labelled with 10-plex TMT Isobaric Label Reagents (Thermo Fisher Scientific, Cat # 90113) following the manufacturer’s instructions. The reactions were quenched using 5µL of 5% hydroxylamine for 30 minutes. Nuclear proteins from astrocytes or neurons for NS and SD groups (3 biological replicates for each) were labeled with the six TMT reagents within a TMT 10-plex reagent set, while the 7-10th reagent was used for unrelated samples. The application of multiplexed TMT reagents allowed comparison of NS and SD samples within the same MS run, which eliminated the possibility of run-to-run (or batch) variations.

### Nano-LC Mass Spectrometry

TMT-labelled peptides were fractionated using an Ultimate 3000 nano-LC system in line with an Orbitrap Fusion Lumos mass spectrometer (Thermo Scientific). In brief, peptides in 1% (vol/vol) formic acid were injected into an Acclaim PepMap C18 nano-trap column (Thermo Scientific). After washing with 0.5% (vol/vol) acetonitrile 0.1% (vol/vol) formic acid peptides were resolved on a 250 mm × 75 μm Acclaim PepMap C18 reverse-phase analytical column (Thermo Scientific) over a 150 min organic gradient, using 7 gradient segments (1-6% solvent B over 1 min., 6-15% B over 58min., 15-32%B over 58min., 32-40%B over 5min., 40-90%B over 1min., held at 90%B for 6min and then reduced to 1%B over 1min.) with a flow rate of 300nL min^−1^. Solvent A was 0.1% formic acid and Solvent B was aqueous 80% acetonitrile in 0.1% formic acid. Each sample was injected three times to improve proteome coverage. Peptides were ionized by nano-electrospray ionization at 2.0kV using a stainless-steel emitter with an internal diameter of 30 μm (Thermo Scientific) and a capillary temperature of 275°C. All spectra were acquired using an Orbitrap Fusion Lumos mass spectrometer controlled by Xcalibur 4.1 software (Thermo Scientific) and operated in data-dependent acquisition mode using an SPS-MS3 workflow. FTMS1 spectra were collected over the range of 375–1500 m/z at a resolution of 120,000, with an automatic gain control (AGC) target of 200,000 and a max injection time of 50 ms. Precursors were filtered with an intensity threshold of 5000, according to charge state (to include charge states 2-7) and with monoisotopic peak determination set to Peptide. Previously interrogated precursors were excluded using a dynamic window (60s +/-10ppm). For FTMS3 analysis, the Orbitrap was operated at 50,000 resolutions with an AGC target of 50,000 and a max injection time of 105 ms. Precursors were fragmented by high energy collision dissociation (HCD) at a normalised collision energy of 60% to ensure maximal TMT reporter ion yield. Synchronous Precursor Selection (SPS) was enabled to include up to 5 MS2 fragment ions in the FTMS3 scan.

### Database search and statistical analysis of quantitative proteomics data

Quantitative nuclear proteomics raw data files were analyzed using the MaxQuant computational platform (*version* 2.1.3.0) with the Andromeda search engine ^65^. MS2/MS3 spectra were searched against UniProt database specifying *Mus musculus* (Mouse) taxonomy (Organism ID: 10090). All searches were performed using “Reporter ion MS2/MS3” with “10-plex TMT” as isobaric labels with a static modification for cysteine alkylation (carbamidomethylation), and oxidation of methionine (M) and protein N-terminal acetylation as the variable modifications. Trypsin digestion with maximum two missed cleavages, minimum peptide length of seven amino acids, precursor ion mass tolerance of 5 ppm, and fragment mass tolerance of 0.02 Da were specified in all analyses. The false discovery rate (FDR) was specified at 0.01 for peptide spectrum match (PSM), protein, and site decoy fraction. TMT signals were corrected for isotope impurities based on the manufacturer’s instructions. Subsequent processing and statistical analysis of quantitative proteomics datasets were performed using the Perseus workstation (*version* 2.0.6.0) ^66^. During data processing, reverse and contaminant database hits and candidates identified only by site were filtered out.

For differential quantitative proteomics analyses, categorical annotation (NS/SD) was applied to group reporter ion intensities, values were Log_2_ transformed and were normalized by “subtract mean (column-wise)” in each TMT reporter ion channel. Protein groups were filtered for valid values (at least 80% in each group). Unpaired t-tests followed by false discovery rate (FDR) analysis were performed to compare NS and SD groups.

### Volcano plots and heatmaps

We used volcano plots and heatmaps to visualize gene and protein expression changes for comparisons of NS and SD mice. The unpaired *t*-tests followed by FDR analysis were performed to test for significant differences in gene expression and protein abundance between NS and SD in astrocytes and neurons. A threshold of 1.3-fold changes in expression and abundance is considered to be defined as downregulation (log_2_(SD/NS) < -0.4, FDR < 0.05) and upregulation (log_2_(SD/NS) > 0.4, FDR < 0.05).

### Gene ontology (GO) analysis

Significantly and differentially expressed transcripts and proteins were subjected to GO analysis. We performed an overrepresentation analysis ^67^ that was implemented in clusterProfiler ^68^. Significantly enriched GO terms and KEEG pathways were visualized as dot plots, cnet plots, and bar plots.

### Motif analysis and predictions of TFs

For TF binding site motifs analysis and prediction of TFs from gene expression data, *Pscan* (*version 1.6*) was used ^22^. The sequence motif analysis was performed by selecting +1000 -0 bp of DNA proximal to TSS and JASPAR database was used to predict TFs. Transcriptomics data were also utilized for TF Co-regulatory Networks analysis by the ChIP-X Enrichment Analysis (*version 3*) (ChEA3) ^28^. The edges between TFs were determined based on evidence from the ChEA3 libraries and are directed according to ChIP-seq data that supports the interaction. The mean rank across the libraries (ChIP-Seq, Coexpression, and Co-occurrence) was used to plot the networks. For the characterization of TFs from nuclear proteins and identification of their target genes, TRRUST version 2 was used ^29^. The mouse TFs and their co-expressed gene database were used. The significantly downregulation (log_2_(SD/NS) < -0.4, FDR < 0.05) and upregulation (log_2_(SD/NS) > 0.4, FDR < 0.05) genes and proteins with 1.3-fold change thresholds were used for these analyses.

### Statistical methods

All experimental subjects are biological replicates. R software was used to perform statistical tests and data visualization. The unpaired t-tests were performed to compare NS and SD groups. The Benjamini-Hochberg procedure was used to calculate the FDR. For transcriptomics, proteomics, and TFs analyses we used FDR < 0.05

## Acknowledgments

A.B.R. acknowledges funding from the Perelman School of Medicine, the University of Pennsylvania, and the Institute for Translational Medicine and Therapeutics (ITMAT) at the University of Pennsylvania. This work was supported also by NIH DP1DK126167 and R01GM139211 (A.B.R.).

## Author contributions

A.B.R., P.K.J., and U.K.V. conceived and designed the experiments. P.K.J. performed mouse sleep deprivation experiments and tissue dissections. P.K.J. and U.K.V. performed single-cell isolation and astrocyte and neuron separation from single-cell suspensions. U.K.V. performed library preparation and quantitative proteomics. U.K.V. and P.K.J. analyzed transcriptomics data. P.K.J. analyzed proteomics data. A.B.R. supervised the entire study and secured funding. The manuscript was written by P.K.J. and A.B.R. with input from U.K.V. All authors agreed on the interpretation of data and approved the final version of the manuscript.

